## Supplementary Information for "Functional metabolomics of the human scalp: A metabolic niche for *Staphylococcus epidermidis*"

### Supplementary Table

**Supplementary Table S1.** Distribution of annotations for each dataset.

| | C18+ | C18− | HILIC+ | HILIC− |
|---|---|---|---|---|
| **Total number of features** | **11,981** | **5,473** | **20,156** | **6,957** |
| Annotated features or in network (GNPS and SIRIUS) | 57% | 95% | 28% | 57% |
| In molecular network (GNPS) | 27% | 53% | 16% | 27% |
| Annotated features (GNPS and SIRIUS) | 54% | 90% | 24% | 54% |
| Mol. formula (SIRIUS) | 53% | 90% | 23% | 53% |
| Mol. formula (ZodScore > 0.7) (SIRIUS) | 32% | 66% | 16% | 32% |
| Chemical class (SIRIUS) | 5% | 52% | 16% | 5% |
| Putative structure (SIRIUS) | 3% | 29% | 11% | 3% |
| Spectral library match (GNPS) | 4% | 2% | 3% | 4% |
| Spectral library match analogue mode (GNPS) | 9% | 9% | 8% | 9% |

### Supplementary Figures

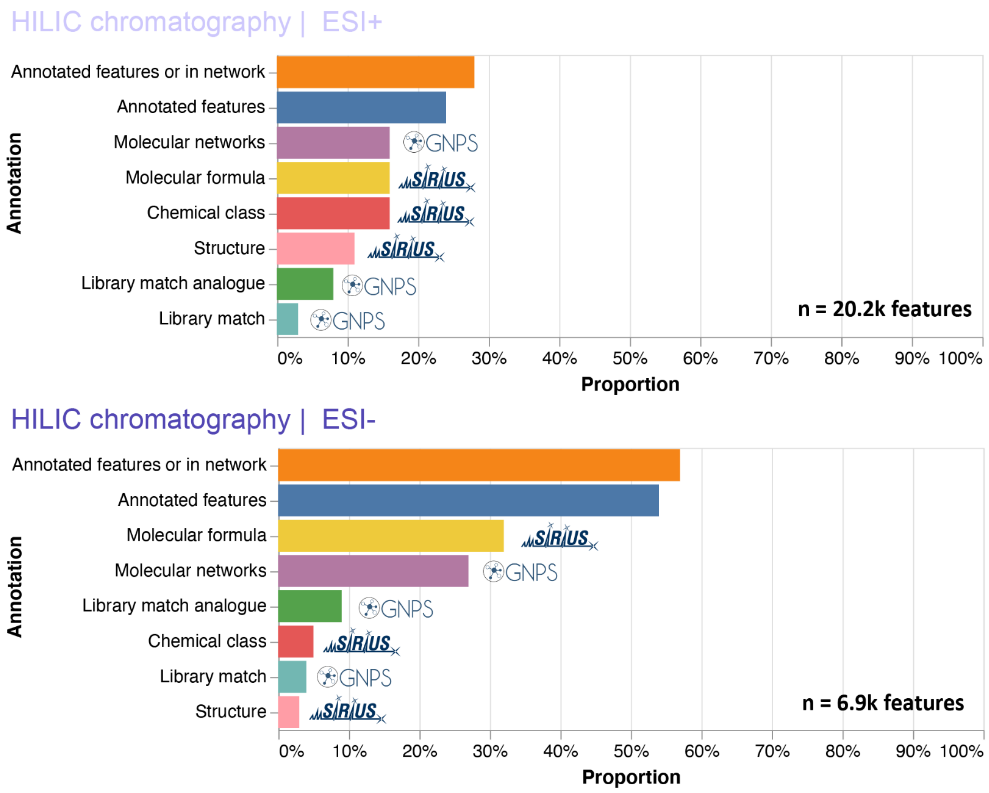


**Supplementary Figure S1.** The proportion of LC-MS/MS features annotated by GNPS or SIRIUS in the HILIC datasets (top, positive ionization mode; bottom, negative ionization mode).

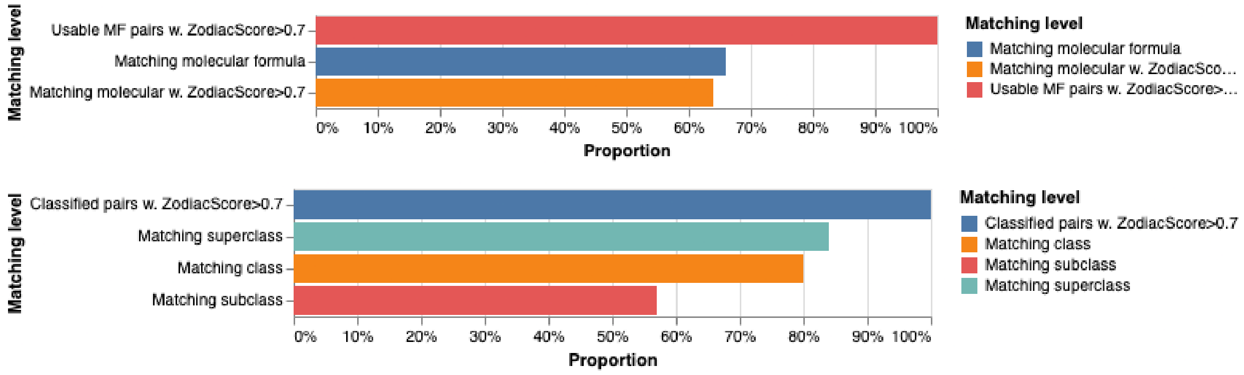


**Supplementary Figure S2.** Evaluating the performance of SIRIUS computational annotation by comparing with usable spectral library matches (n= 44) for the C18+ dataset. The top panel shows the results for molecular formula results. The lower panel shows the results for chemical class annotation.

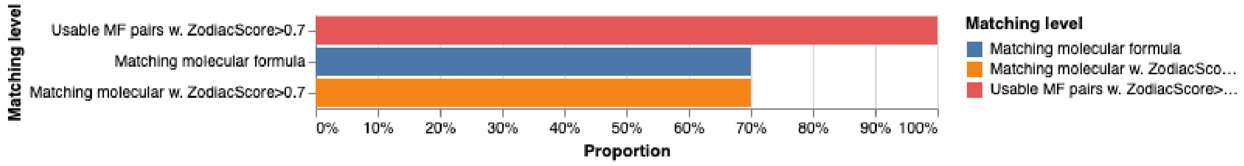

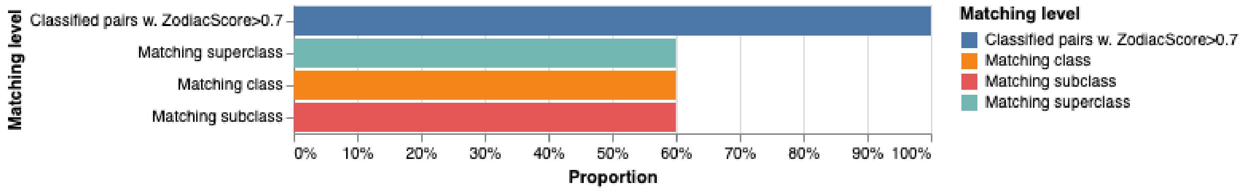


**Supplementary Figure S3.** Evaluating the performance of SIRIUS computational annotation by comparing with usable spectral library matches (n = 10) for the C18− dataset. The top panel shows the results for molecular formula results. The lower panel shows the results for chemical class annotation.

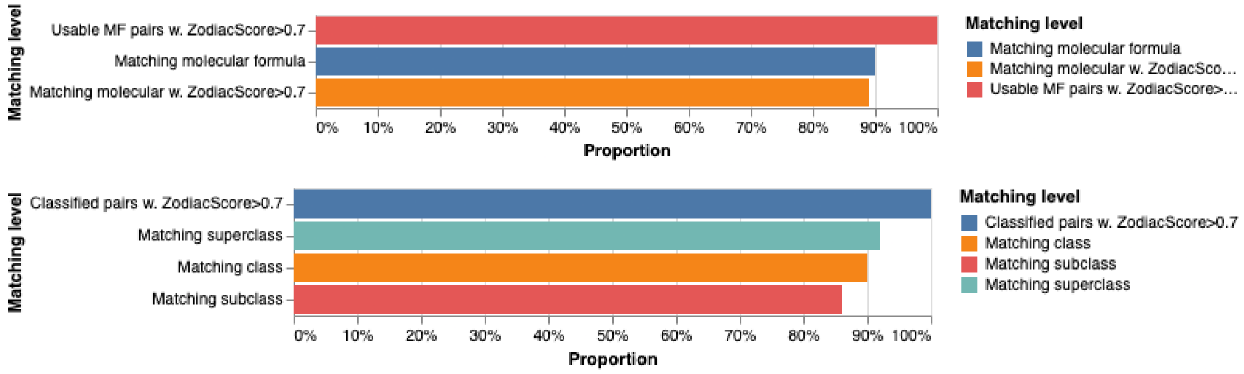


**Supplementary Figure S4.** Evaluating the performance of SIRIUS computational annotation by comparing with usable spectral library matches (n =125) for the HILIC+ dataset. The top panel shows the results for molecular formula results. The lower panel shows the results for chemical class annotation.

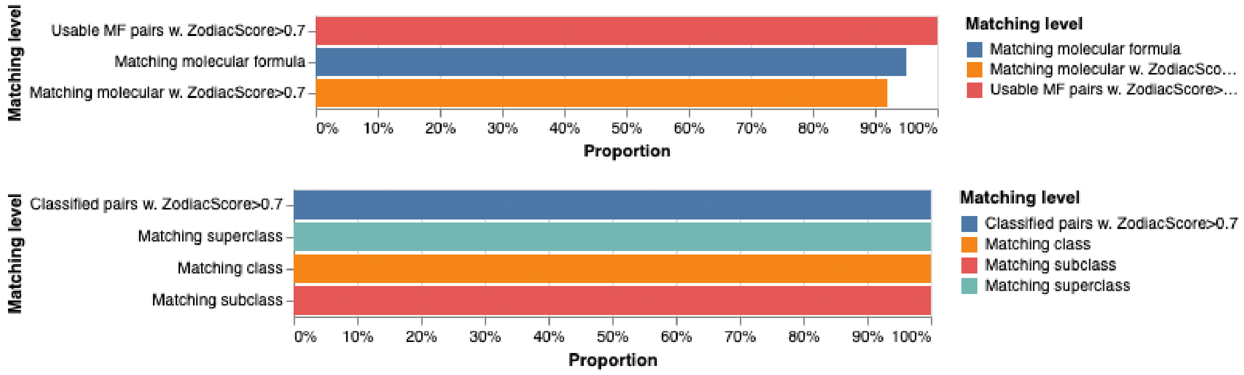


**Supplementary Figure S5.** Evaluating the performance of SIRIUS computational annotation by comparing with usable spectral library matches (n = 59) for the HILIC− dataset. The top panel shows the results for molecular formula results. The lower panel shows the results for chemical class annotation.

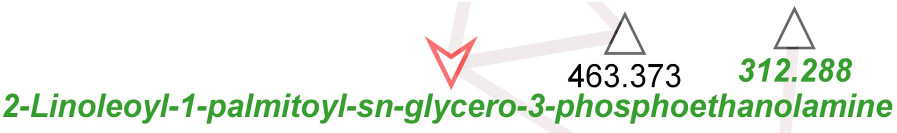


m/z = 575.5076
Molecular formula (SIRIUS): —
GNPS library annotation: 2-Linoleoyl-1-palmitoyl-sn-glycero-3-phosphoethanolamine
Phyla with microbeMASST hits ($p < 0.01$, Fisher's exact test*): Zoopagomycota
Tree of all microbeMASST hits:

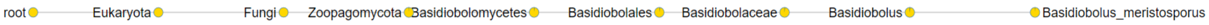


**Supplementary Figure S6.** Detailed annotation and microbMASST results for the feature X9421045

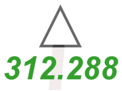


m/z = 312.288
Molecular formula (SIRIUS): $C_{19}H_{39}NO_3$
GNPS library annotation (analog mode**): N-cis-Hexadec-9-enoyl-L-homoserine lactone
Phyla with microbeMASST hits ($p < 0.01$, Fisher's exact test*): Bacteroidetes
Tree of all microbeMASST hits:

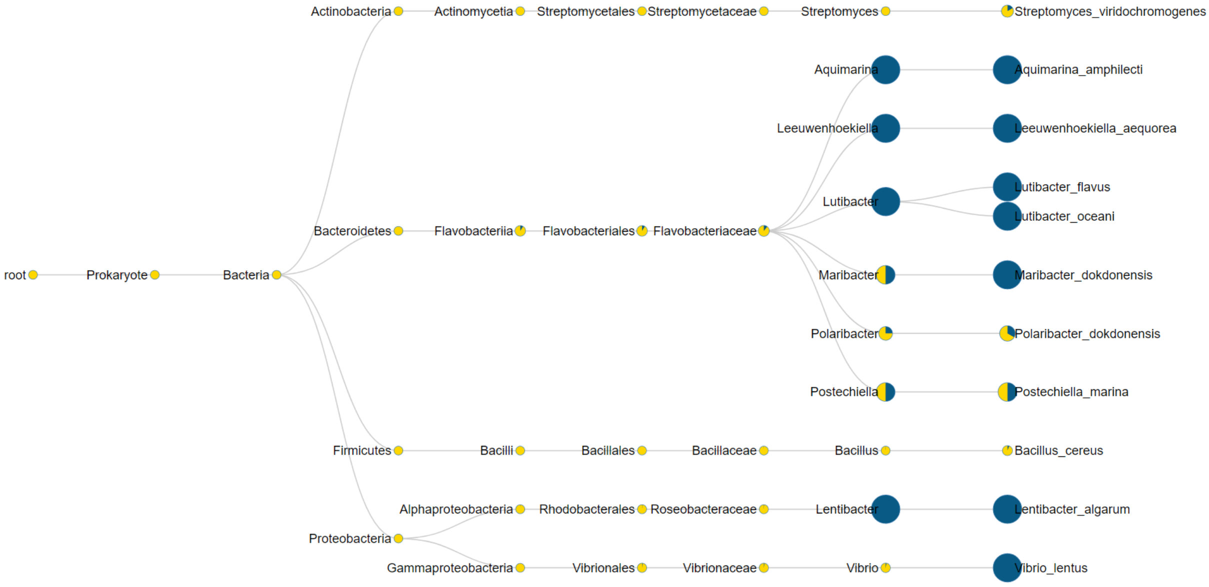


**Supplementary Figure S7.** Detailed annotation and microbMASST results for feature X9421657

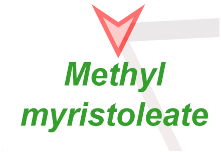


m/z = 241.215
Molecular formula (SIRIUS): C15H28O2
GNPS library annotation: Methyl myristoleate
Phyla with microbeMASST hits ($p < 0.01$, Fisher's exact test*): Chordata
Tree of all microbeMASST hits:

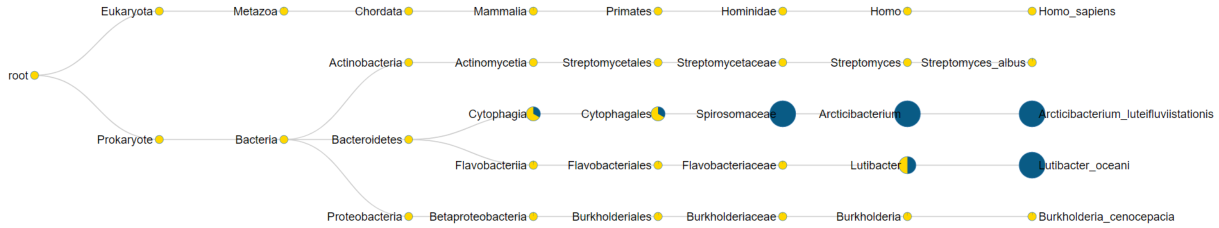


**Supplementary Figure S8.** Detailed annotation and microbMASST results for the feature X9423263

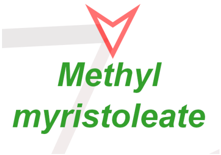


m/z = 241.2149
Molecular formula (SIRIUS): C15H28O2
GNPS library annotation: Methyl myristoleate
Phyla with microbeMASST hits ($p < 0.01$, Fisher's exact test*): Chordata
Tree of all microbeMASST hits:

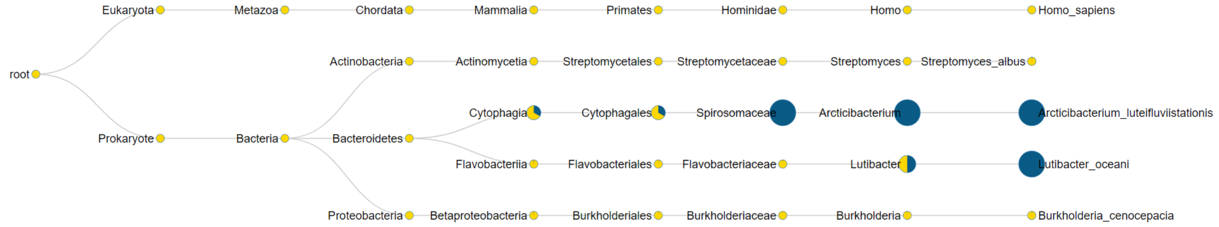


**Supplementary Figure S9.** Detailed annotation and microbMASST results for the feature X9423629

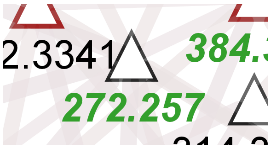


m/z = 272.257

Molecular formula (SIRIUS): C16H30O2

GNPS library annotation (analog mode**): Methyl myristoleate

Phyla with microbeMASST hits ($p < 0.01$, Fisher's exact test*): Ascomycota

Tree of all microbeMASST hits:

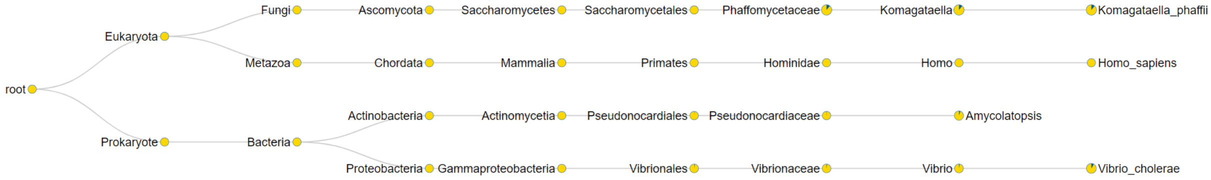


**Supplementary Figure S10.** Detailed annotation and microbMASST results for the feature X9426846

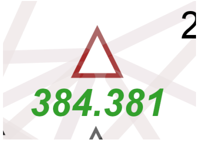


m/z = 384.381

Molecular formula (SIRIUS): C24H46O2

GNPS library annotation (analog mode**): Monoerucin

Phyla with microbeMASST hits ($p < 0.01$, Fisher's exact test*): Ascomycota, Actinobacteria, Firmicutes, Bacteroidetes, Proteobacteria

Tree of all microbeMASST hits:

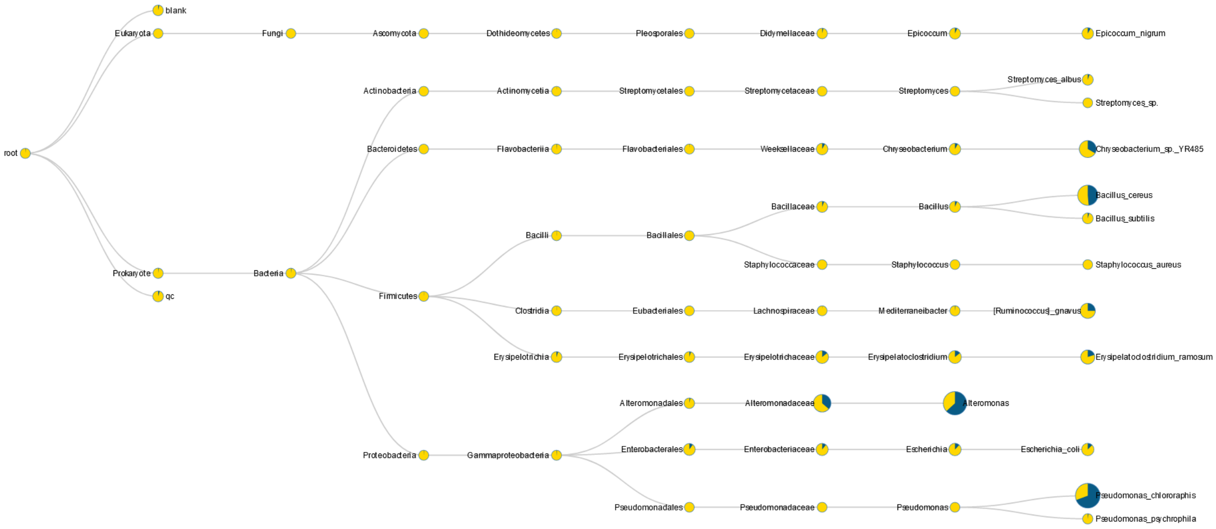


**Supplementary Figure S11.** Detailed annotation and microbMASST results for the feature X9404581

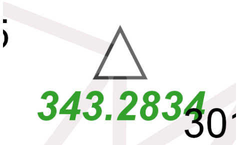


m/z = 343.2834
Molecular formula (SIRIUS): C20H38O4
GNPS library annotation (analog mode**): Monoelaidin
Phyla with microbeMASST hits ($p < 0.01$, Fisher’s exact test*): Actinobacteria, Proteobacteria
Tree of all microbeMASST hits:

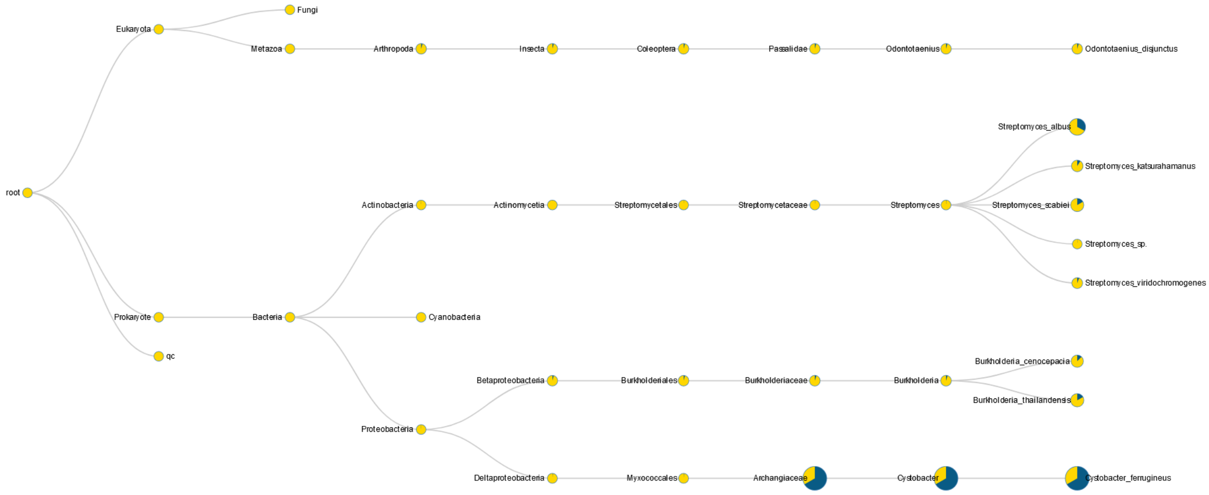


**Supplementary Figure S12.** Detailed annotation and microbMASST results for the feature X9407909

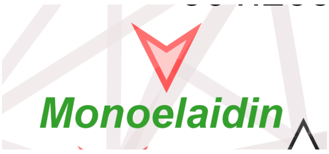


m/z = 357.2983
Molecular formula (SIRIUS): C21H40O4
GNPS library annotation: Monoelaidin
Phyla with microbeMASST hits ($p < 0.01$, Fisher's exact test*): Chordata, Zoopagomycota, Basidiomycota, Ascomycota, Actinobacteria, Firmicutes, Spirochaetes, Proteobacteria
Tree of all microbeMASST hits:

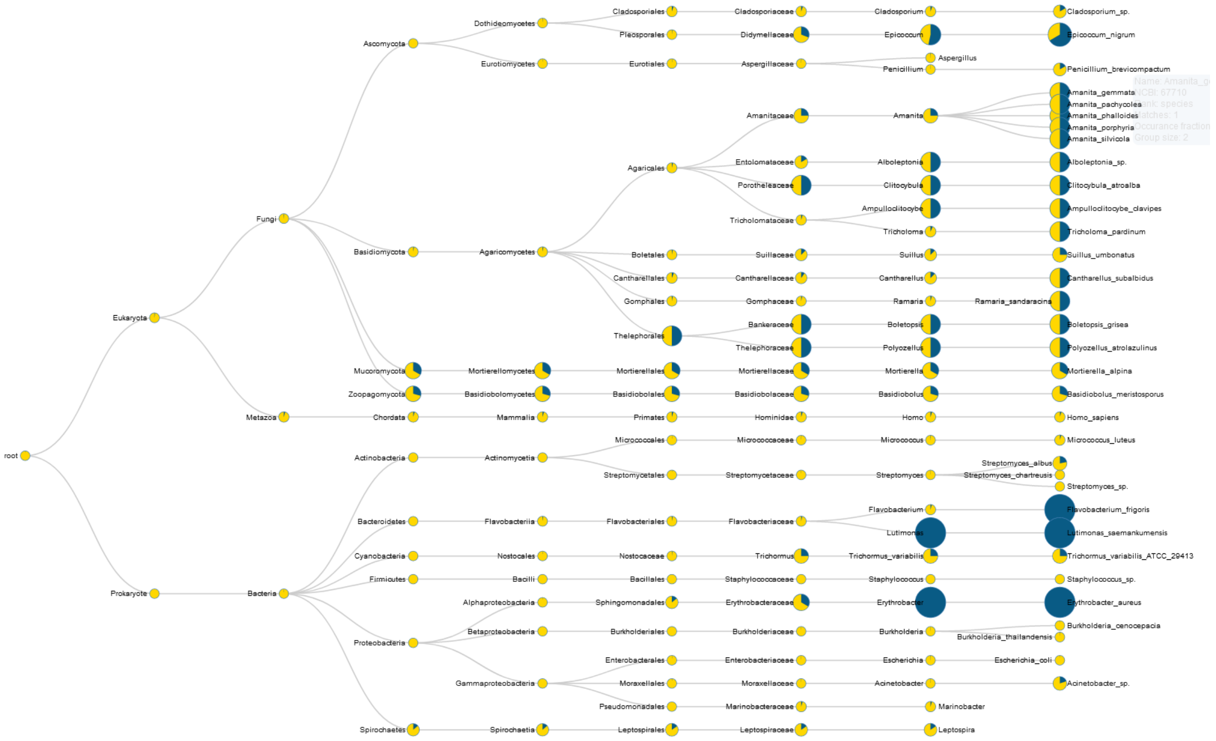


**Supplementary Figure S13.** Detailed annotation and microbMASST results for the feature X9400316

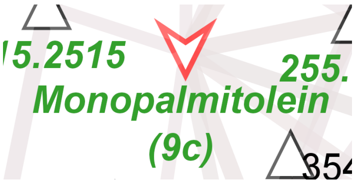


m/z = 329.2673
Molecular formula (SIRIUS): C19H36O4
GNPS library annotation: Monopalmitolein (9c)
Phyla with microbeMASST hits (p < 0.01, Fisher's exact test*): Arthropoda, Cyanobacteria, Firmicutes, Spirochaetes, Proteobacteria
Tree of all microbeMASST hits:

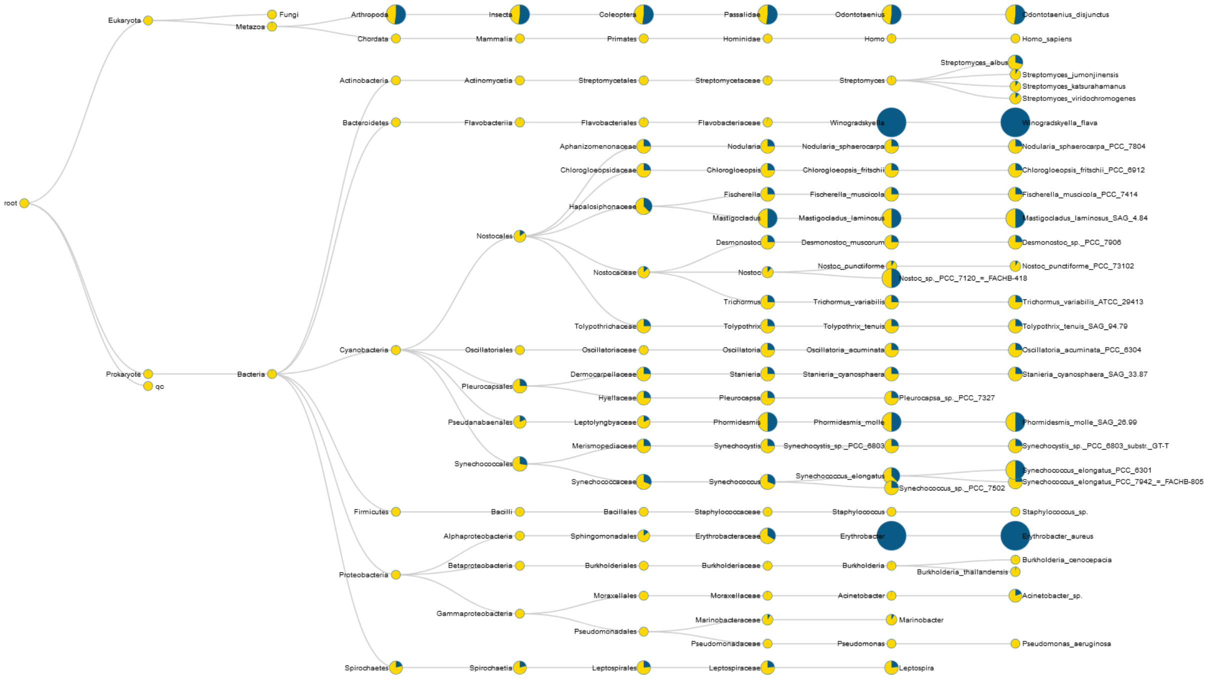


**Supplementary Figure S14.** Detailed annotation and microbMASST results for the feature X9400257

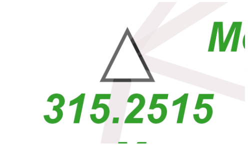


m/z = 315.2515
Molecular formula (SIRIUS): C17H32N4
GNPS library annotation (analog mode**): Monoolein
Phyla with microbeMASST hits ($p < 0.01$, Fisher’s exact test*): Arthropoda
Tree of all microbeMASST hits:

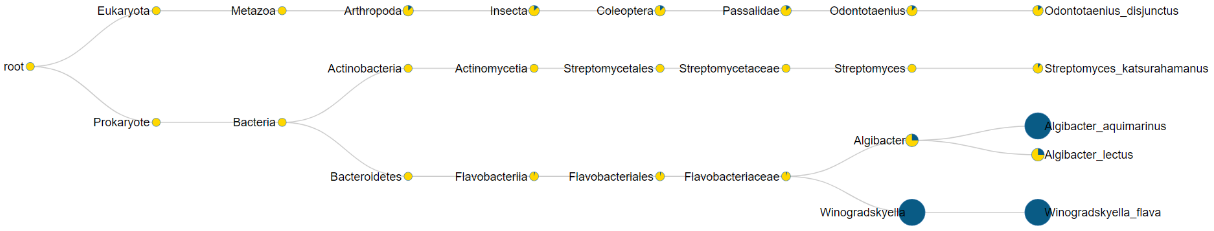


**Supplementary Figure S15.** Detailed annotation and microbMASST results for the feature X9402417

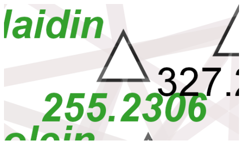


m/z = 255.2306
Molecular formula (SIRIUS): C16H30O2
GNPS library annotation (analog mode**): Palmitoleic Acid ethyl ester
Phyla with microbeMASST hits ($p < 0.01$, Fisher’s exact test*): Actinobacteria, Cyanobacteria, Spirochaetes
Tree of all microbeMASST hits:

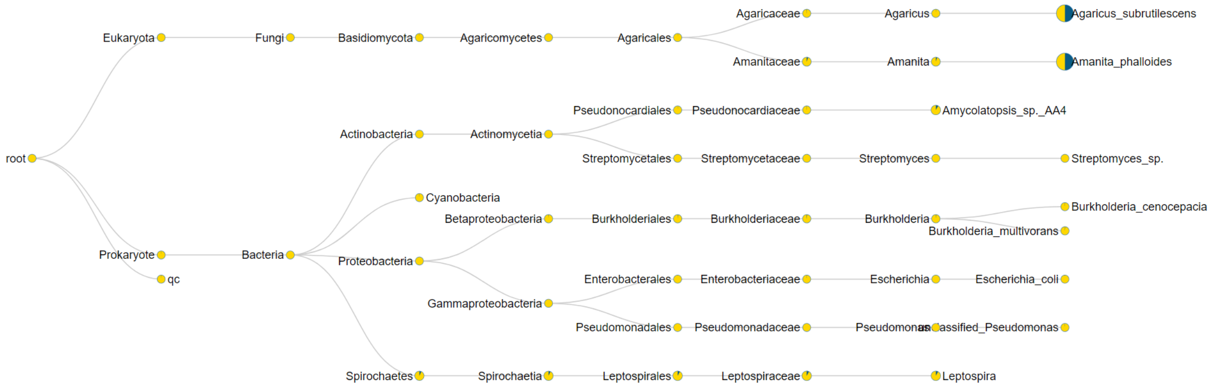


**Supplementary Figure S16.** Detailed annotation and microbMASST results for the feature X9401123

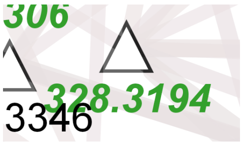


m/z = 328.3194
Molecular formula (SIRIUS): C20H38O2
GNPS library annotation (analog mode**): Oleic acid ethyl ester
Phyla with microbeMASST hits ($p < 0.01$, Fisher's exact test*): Ascomycota, Cyanobacteria
Tree of all microbeMASST hits:

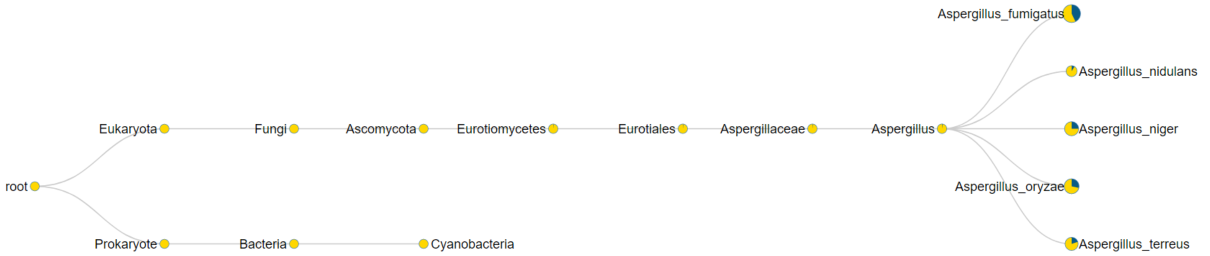


**Supplementary Figure S17.** Detailed annotation and microbMASST results for the feature X9400904

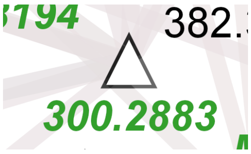


m/z = 300.2883
Molecular formula (SIRIUS): C18H34O2
GNPS library annotation (analog mode**): cis-7-Hexadecenoic acid methyl ester
Phyla with microbeMASST hits ($p < 0.01$, Fisher's exact test*): Cyanobacteria
Tree of all microbeMASST hits:

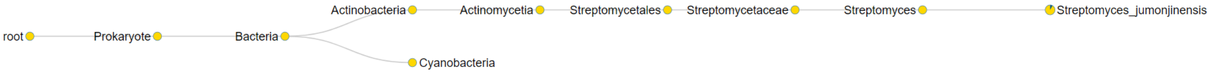


**Supplementary Figure S18.** Detailed annotation and microbMASST results for the feature X9402674

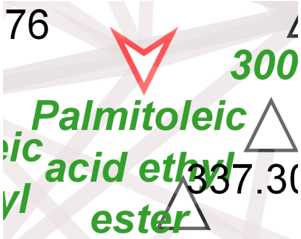


m/z = 283.2625
Molecular formula (SIRIUS): C18H34O2
GNPS library annotation: Palmitoleic acid ethyl ester
Phyla with microbeMASST hits ($p < 0.01$, Fisher's exact test*): Ascomycota, Cyanobacteria, Firmicutes, Proteobacteria
Tree of all microbeMASST hits:

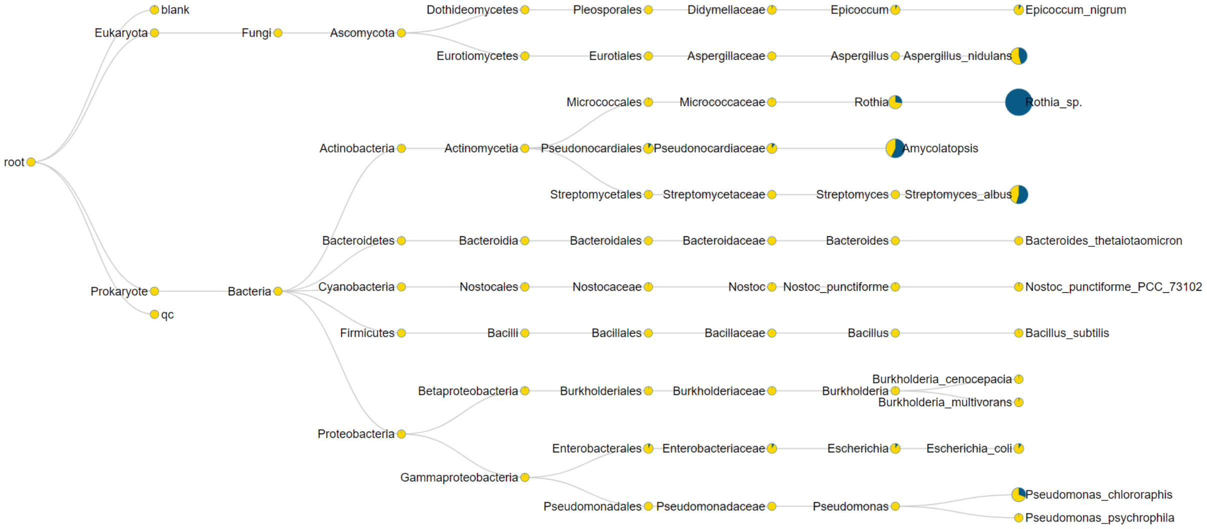


**Supplementary Figure S19.** Detailed annotation and microbMASST results for the feature X9400174

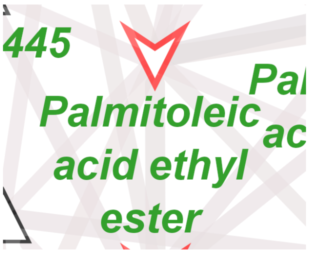


m/z = 283.2614
Molecular formula (SIRIUS): C18H34O2
GNPS library annotation: Palmitoleic acid ethyl ester
Phyla with microbeMASST hits ($p < 0.01$, Fisher's exact test*): Basidiomycota, Ascomycota, Cyanobacteria, Firmicutes, Proteobacteria
Tree of all microbeMASST hits:

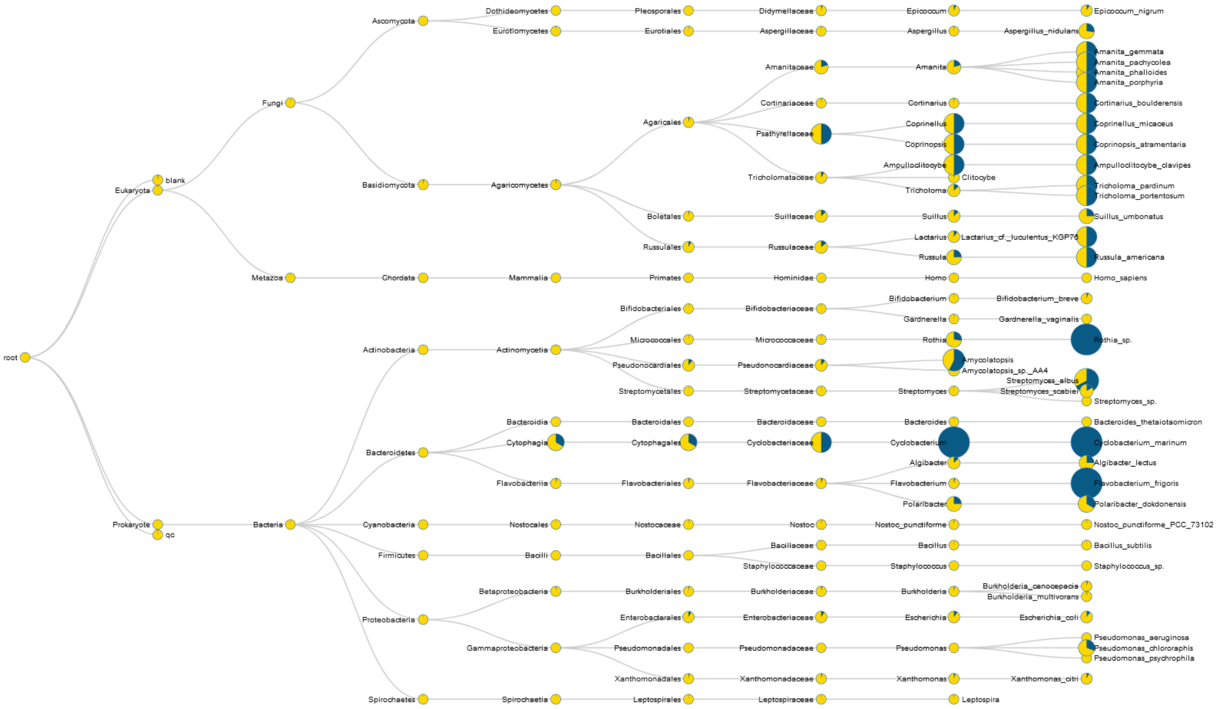


**Supplementary Figure S20.** Detailed annotation and microbMASST results for the feature X9404362

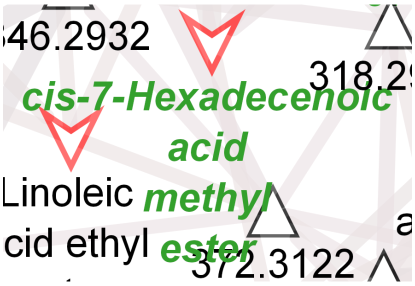

m/z = 269.2461
Molecular formula (SIRIUS): $C_{17}H_{32}O_2$
GNPS library annotation: cis-7-Hexadecenoic acid methyl ester
Phyla with microbeMASST hits ($p < 0.01$, Fisher's exact test*): Ascomycota, Actinobacteria, Firmicutes, Proteobacteria
Tree of all microbeMASST hits:

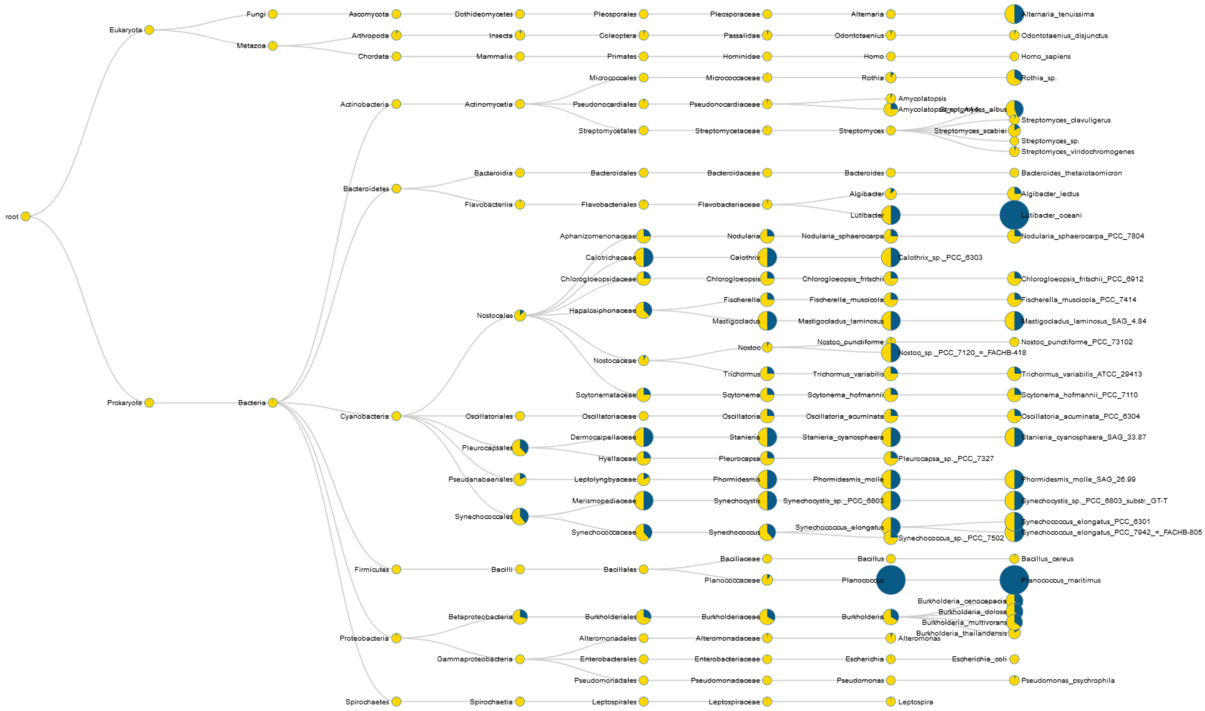

**Supplementary Figure S21.** Detailed annotation and microbMASST results for the feature X9409048

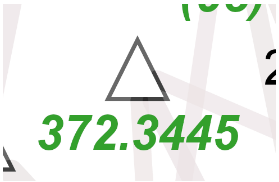


m/z = 372.3445
Molecular formula (SIRIUS): C22H42O3
GNPS library annotation (analog mode**): 1-Linoleoylglycerol
Phyla with microbeMASST hits ($p < 0.01$, Fisher's exact test*): Basidiomycota, Ascomycota
Tree of all microbeMASST hits:

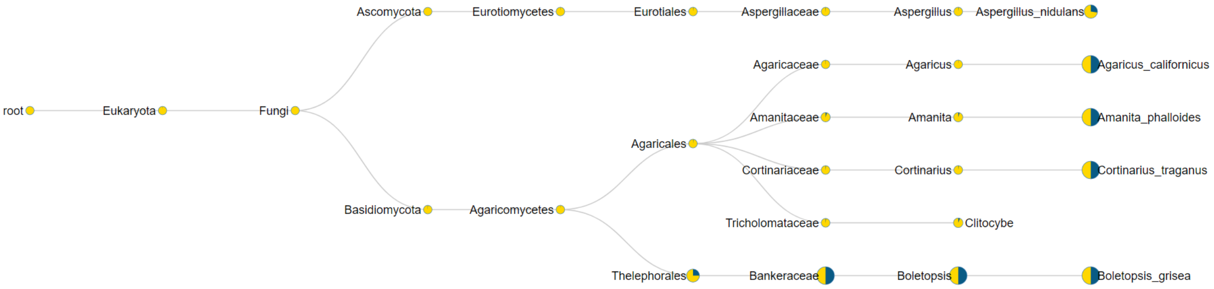


**Supplementary Figure S22.** Detailed annotation and microbMASST results for the feature X9430538

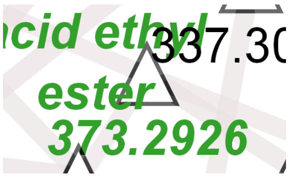


m/z = 373.2926
Molecular formula (SIRIUS): C21H40O5
GNPS library annotation (analog mode**): 2-Linoleoylglycerol
Phyla with microbeMASST hits ($p < 0.01$, Fisher's exact test*): Actinobacteria
Tree of all microbeMASST hits:

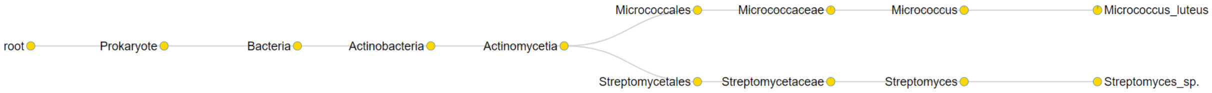


**Supplementary Figure S23.** Detailed annotation and microbMASST results for the feature X9402498

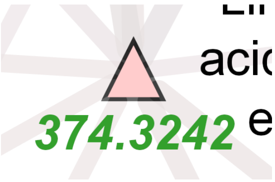


m/z = 374.3242
Molecular formula (SIRIUS): C21H40O4
GNPS library annotation (analog mode**): Monoolein
Phyla with microbeMASST hits (p < 0.01, Fisher's exact test*): Basidiomycota
Tree of all microbeMASST hits:

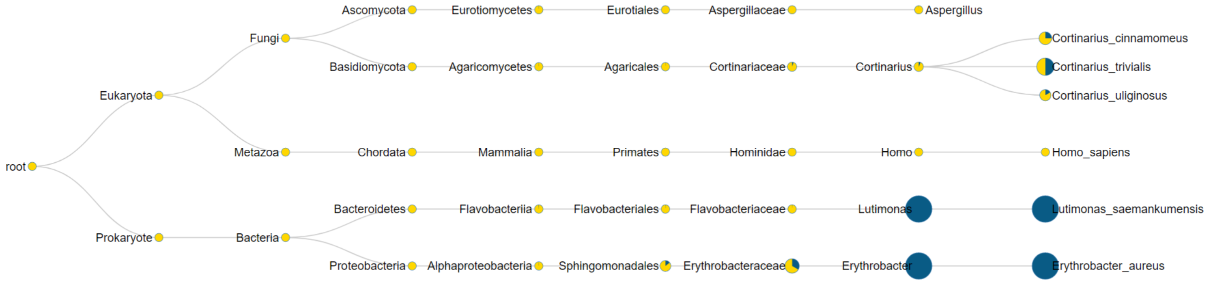


**Supplementary Figure S24.** Detailed annotation and microbMASST results for the feature X9401121

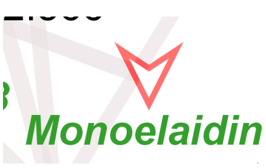


m/z = 339.2871
Molecular formula (SIRIUS): C19H40O3
GNPS library annotation: Monoelaidin
Phyla with microbeMASST hits ($p < 0.01$, Fisher's exact test*): Chordata, Ascomycota, Actinobacteria, Firmicutes, Proteobacteria
Tree of all microbeMASST hits:

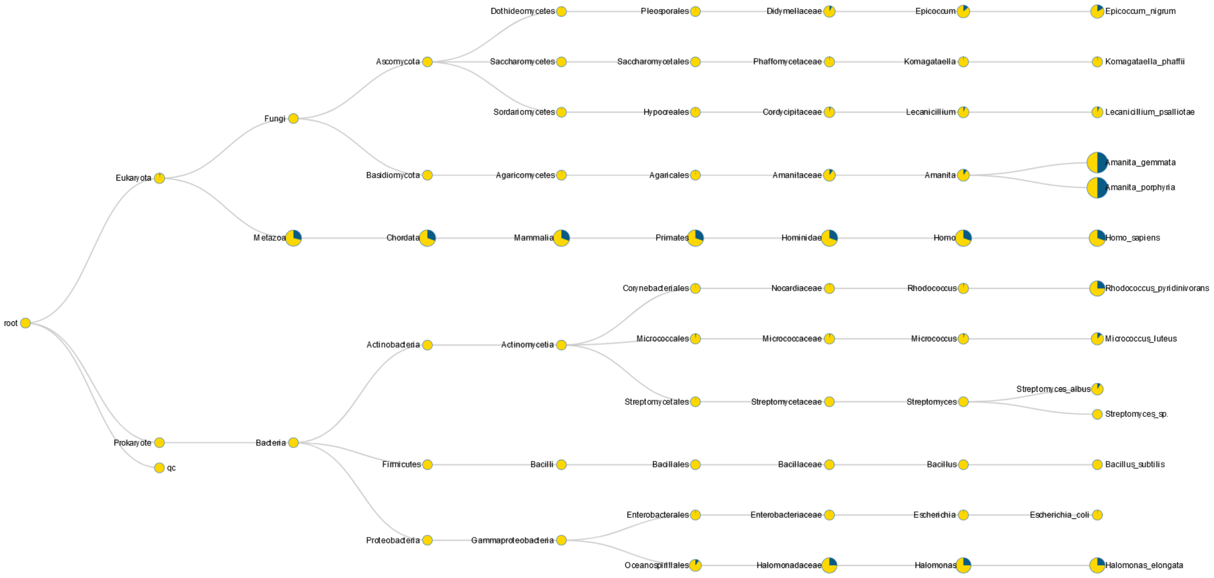


**Supplementary Figure S25.** Detailed annotation and microbMASST results for the feature X9433405

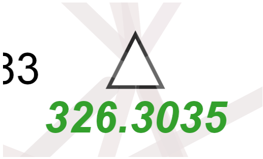


m/z = 326.3035 C20H36O2
Molecular formula (SIRIUS):
GNPS library annotation (analog mode**): Linoleic acid methyl ester
Likely true annotation: Linoleic acid ethyl ester
Phyla with microbeMASST hits ($p < 0.01$, Fisher's exact test*): Proteobacteria
Tree of all microbeMASST hits:

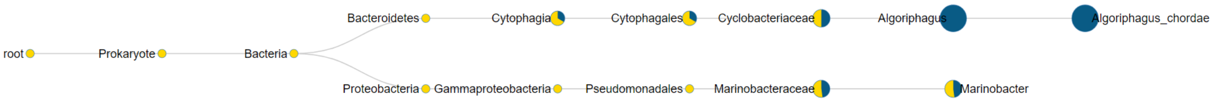


**Supplementary Figure S26.** Detailed annotation and microbMASST results for the feature X9402192

* Significant enrichment of microbeMASST hits in a particular phylum was tested using Fisher’s exact test: The number of microbeMASST hits and the group size of that phylum were compared to the total number of microbeMASST hits across all phyla and the group size of the root node. The null hypothesis was that microbeMASST hits were randomly distributed across all phyla.

** GNPS library annotations in analog mode differ in their precursor m/z from the feature in the metabolomics dataset. As a consequence, the molecular formula predicted by SIRIUS (which will match the precursor m/z) will not match the GNPS library annotation in analog mode. For example, “Supplementary Figure S26. Detailed annotation and microbMASST results for the feature X9402192” has ‘Linoleic acid methyl ester’ as GNPS library annotation in analog mode, which would have the molecular formula C19H34O2. In reality, this feature is likely to be a ‘Linoleic acid ethyl ester’, which would match the molecular formula C20H36O2 predicted by SIRIUS.

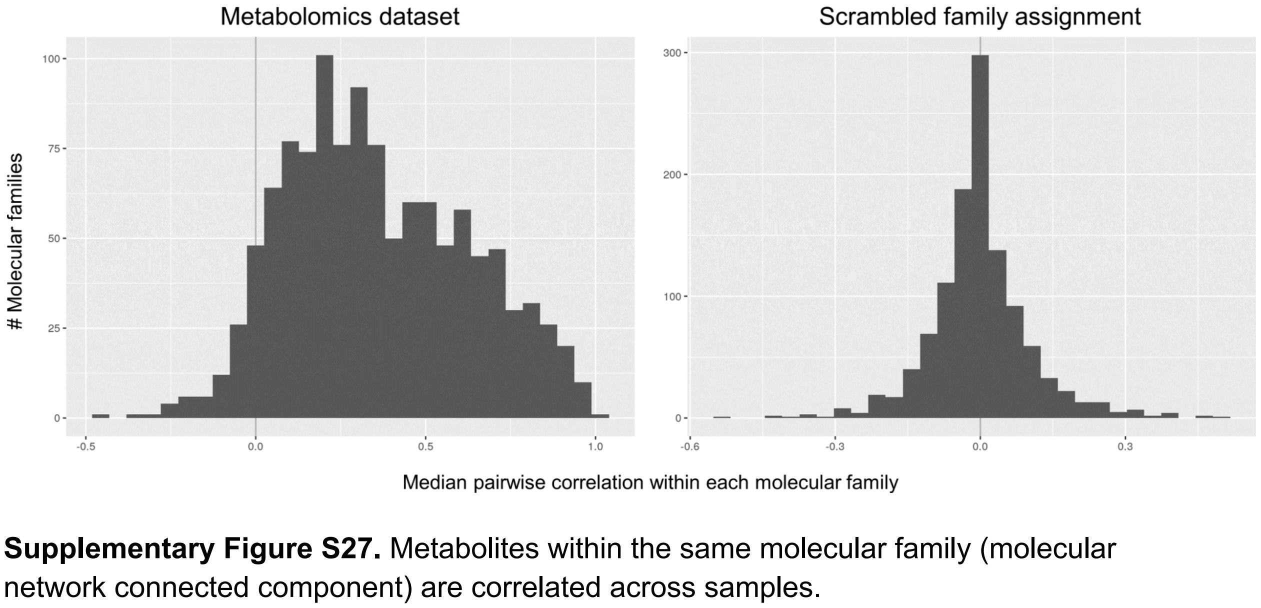


**Supplementary Figure S27.** Metabolites within the same molecular family (molecular network connected component) are correlated across samples.

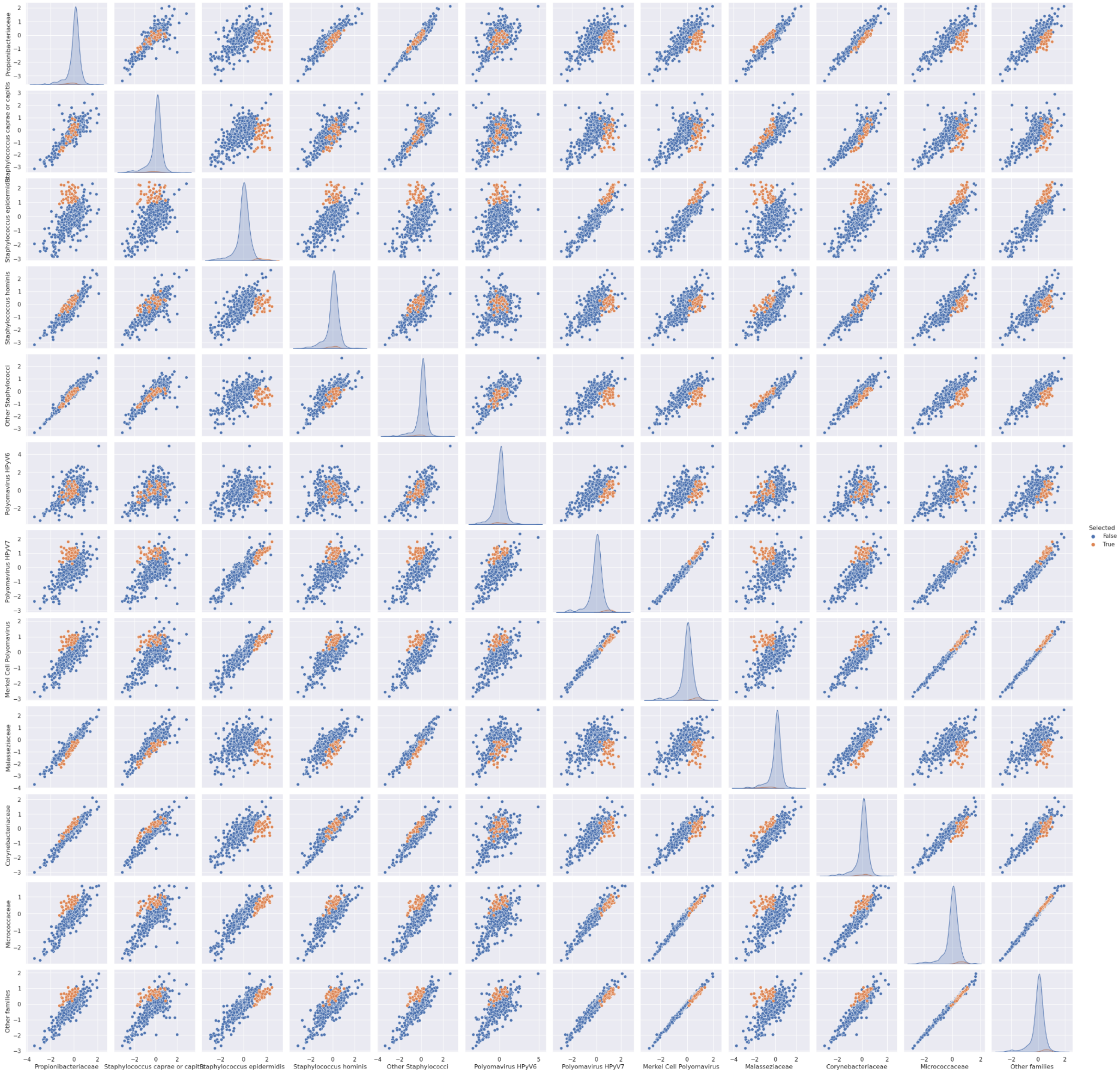


**Supplementary Figure S28.** Pairwise correlations of MMvec log conditional probabilities (first of three independent runs). A group of molecular families showing differential co-occurrence probabilities with *S. epidermidis* as compared to Propionibacteriaceae is highlighted in red.

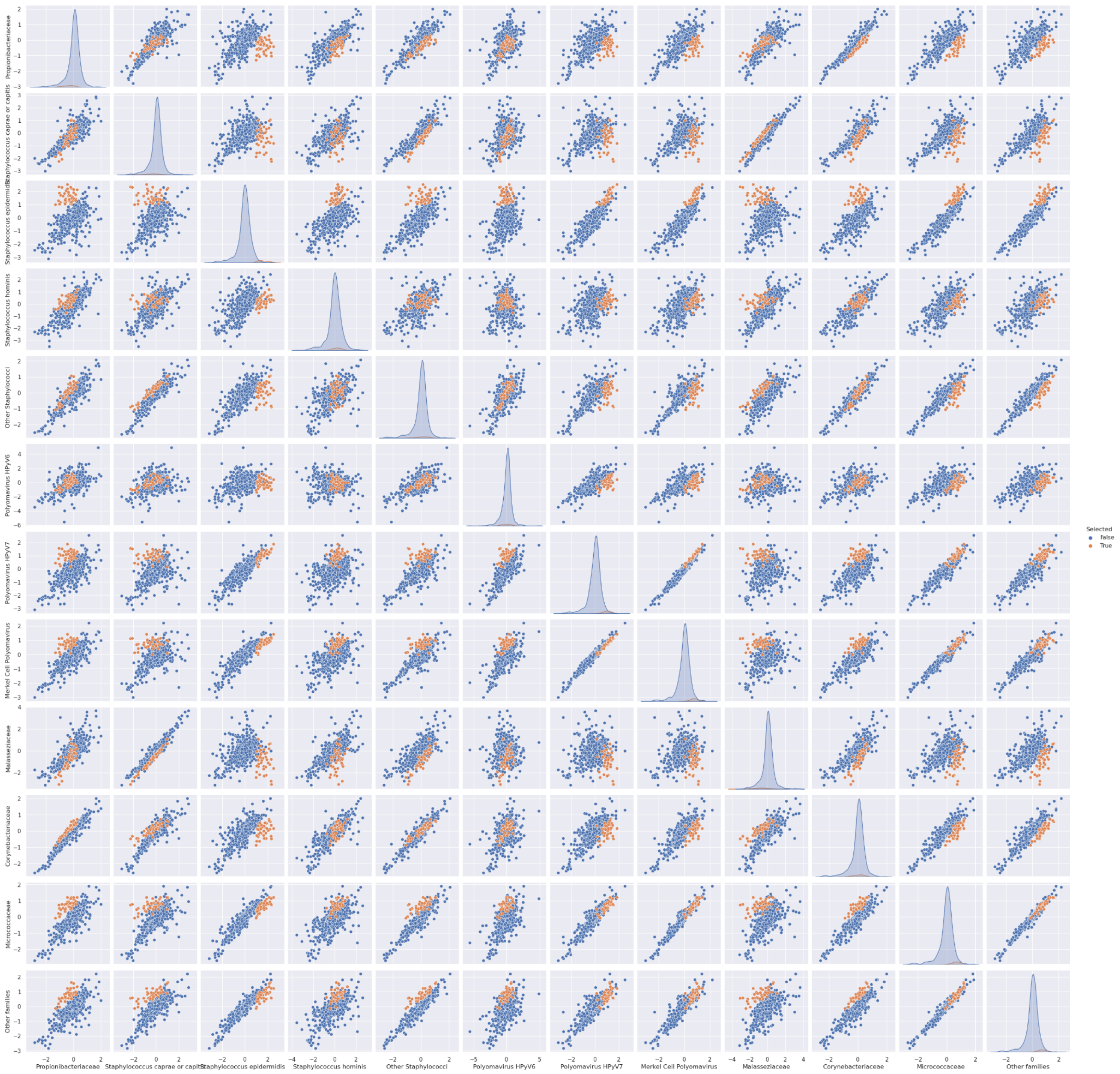

**Supplementary Figure S29.** Pairwise correlations of MMvec log conditional probabilities (second of three independent runs). A group of molecular families showing differential co-occurrence probabilities with *S. epidermidis* as compared to Propionibacteriaceae is highlighted in red.

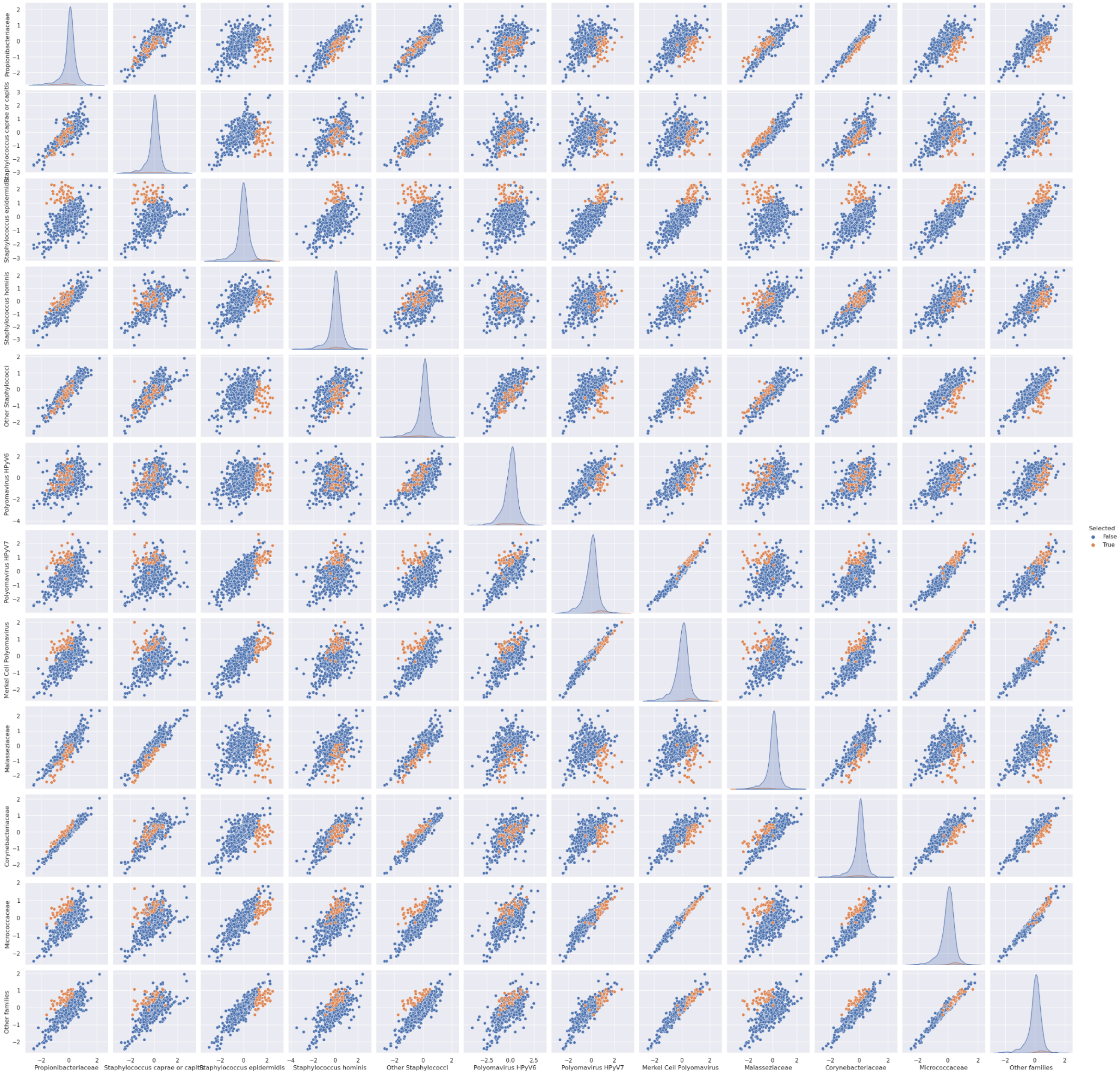


**Supplementary Figure S30.** Pairwise correlations of MMvec log conditional probabilities (third of three independent runs). A group of molecular families showing differential co-occurrence probabilities with *S. epidermidis* as compared to Propionibacteriaceae is highlighted in red.

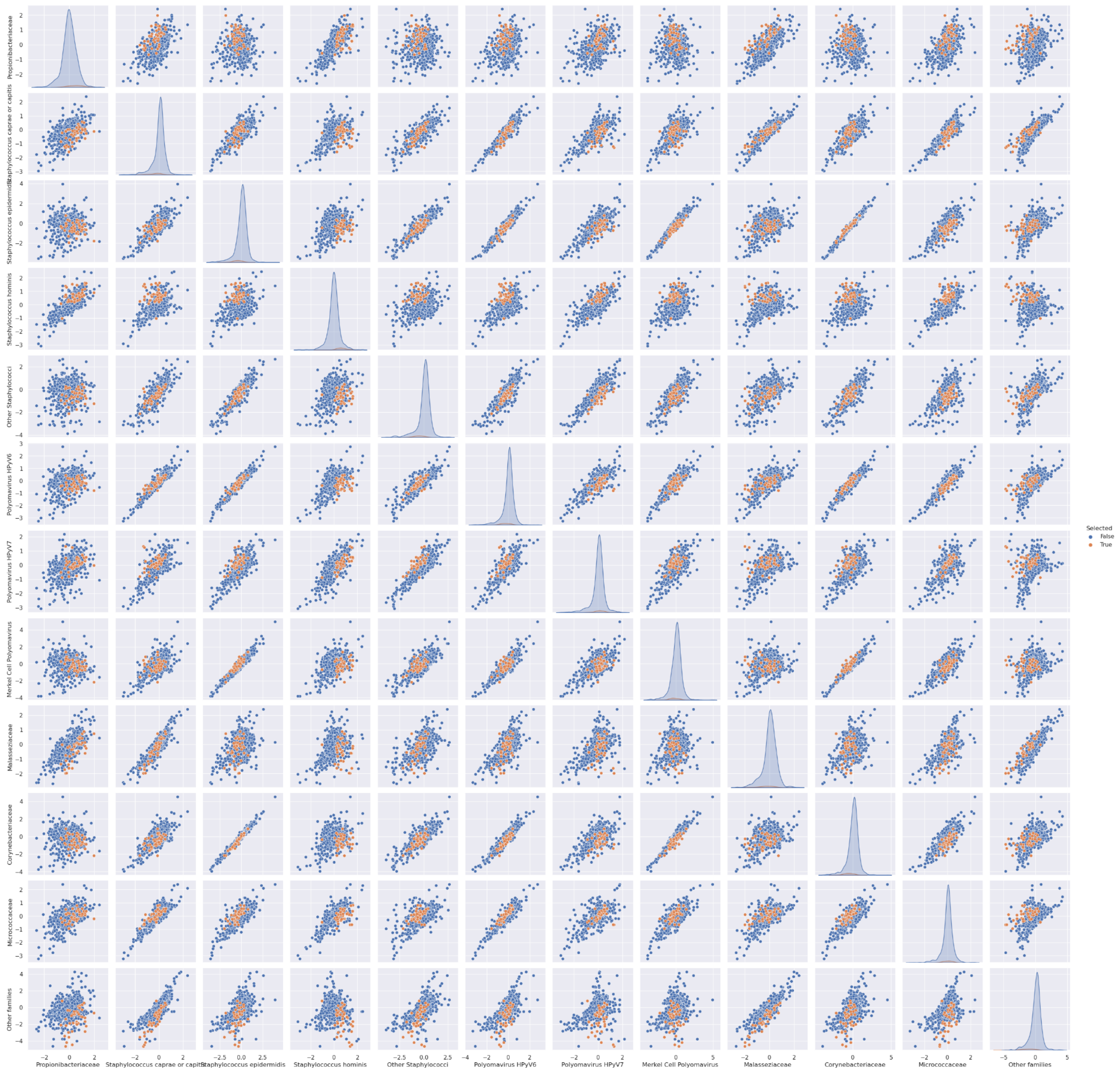


**Supplementary Figure S31.** Pairwise correlations of MMvec log conditional probabilities (scrambled sample identifiers as negative control; first of three independent runs).

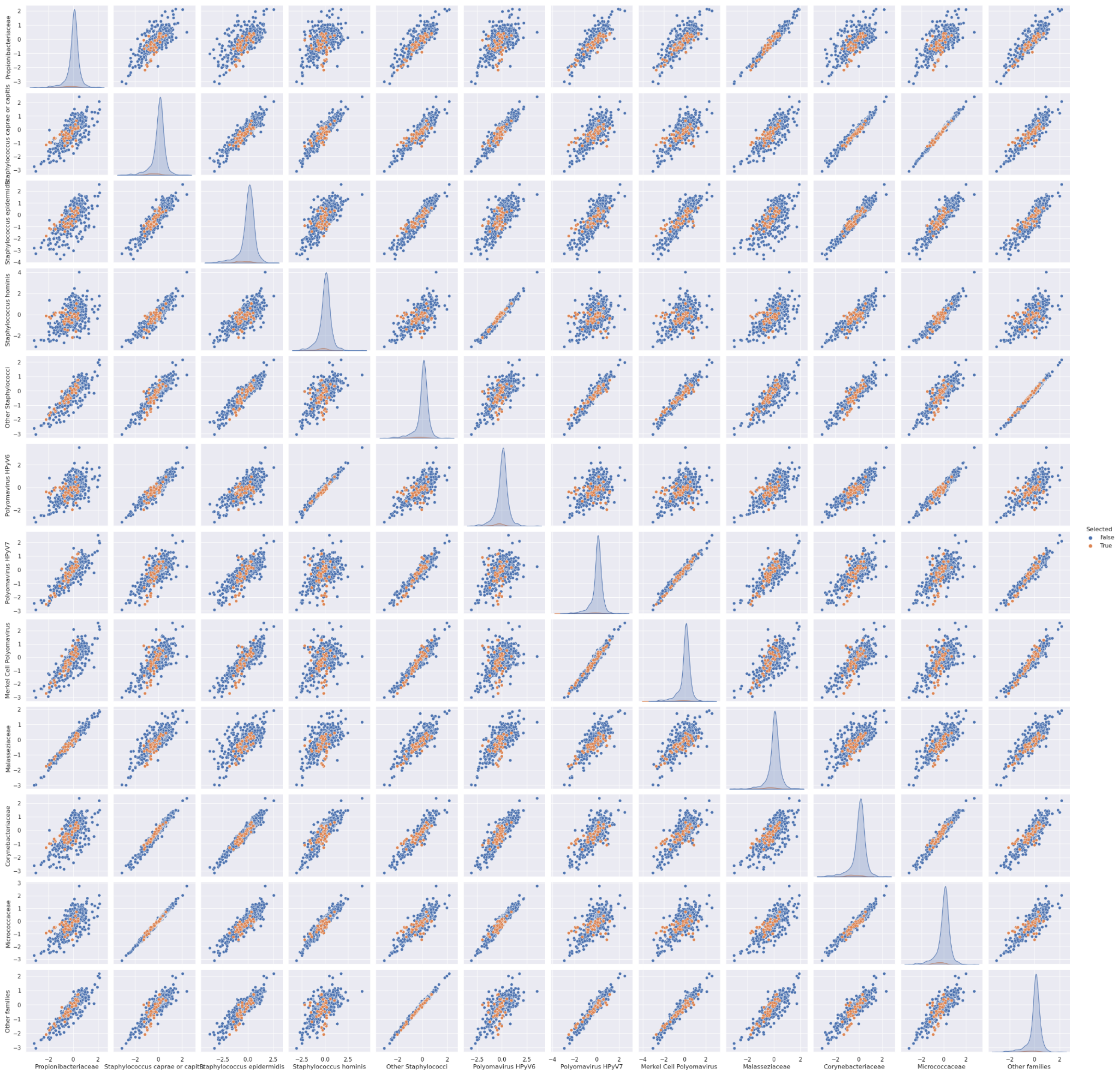


**Supplementary Figure S32.** Pairwise correlations of MMvec log conditional probabilities (scrambled sample identifiers as negative control; second of three independent runs).

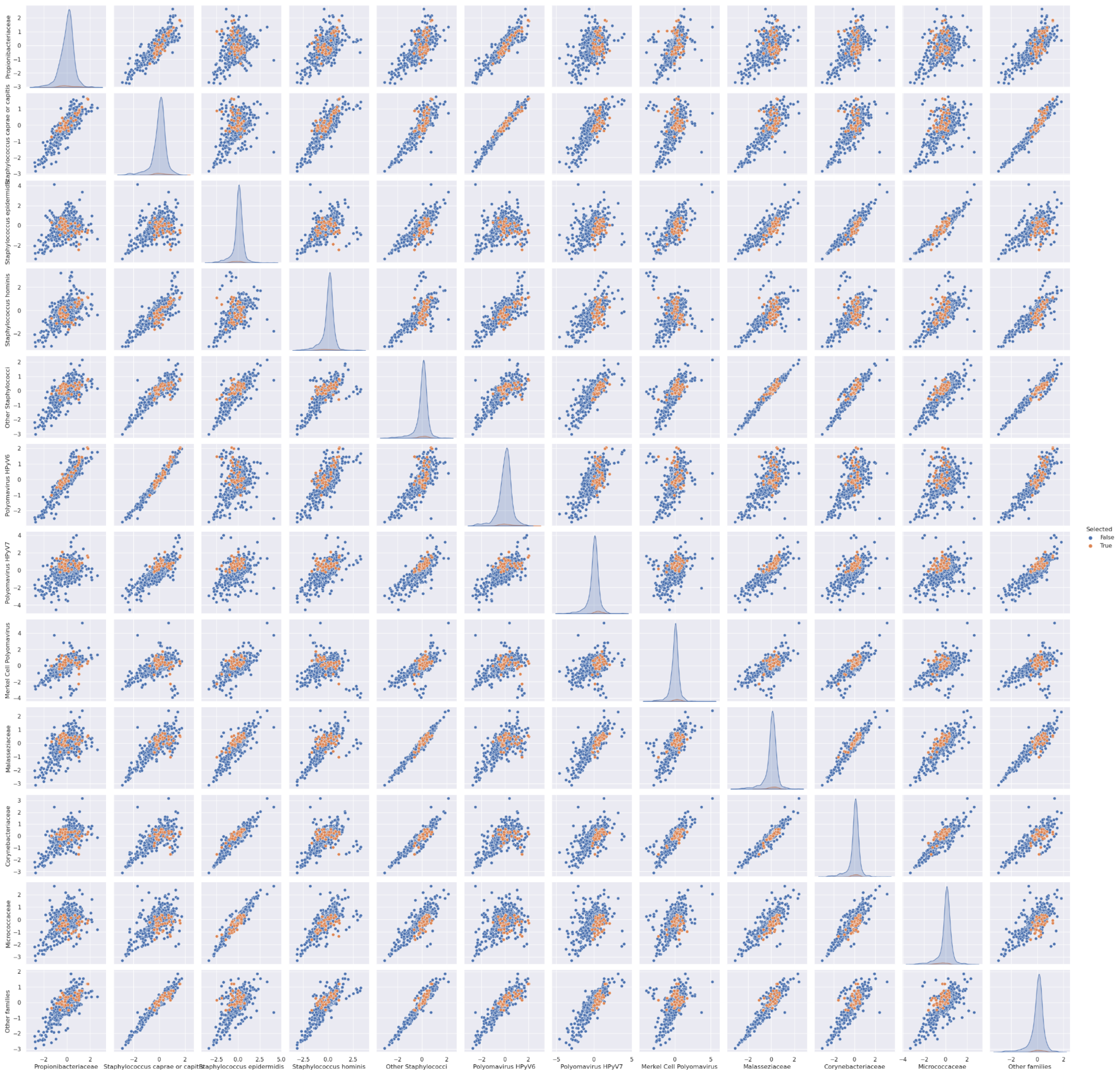


**Supplementary Figure S33.** Pairwise correlations of MMvec log conditional probabilities (scrambled sample identifiers as negative control; third of three independent runs).
